# Beyond seed counts: what two decades of seed testing reveal about European beech in Poland

**DOI:** 10.64898/2026.05.21.726811

**Authors:** Hanna Fuchs, Marcin K. Dyderski, Szymon Jastrzębowski, Ewelina Ratajczak

## Abstract

Forest regeneration depends on a long chain of processes, of which the number of seeds produced is only the first link. Climate-driven changes in seed production and masting in temperate trees are well documented, whereas long-term changes in the quality of the seeds produced have been examined far less often. Using a nationwide operational dataset for European beech (*Fagus sylvatica* L.) from Poland (13,387 seed lots from 380 forest districts, 1996 to 2024; temporal analyses on 5,381 freshly harvested lots from 352 districts, 2004 to 2023), we followed three dimensions of seed quality: seed mass, the proportion of filled seeds (purity) and the viability of filled seeds. The three dimensions were nearly uncorrelated (ρ ≤ 0.11 at the district-year level) and changed over time in different ways. Adjusted annual mean seed mass declined by 1.48 g per year (29.6 g, or about 10%, over 20 years), with the whole distribution shifting downwards uniformly across the country. The median proportion of filled seeds remained stable, but the frequency of lots with fewer than 85% filled seeds rose from about 1% to 14% between 2006 and 2023. The viability of filled seeds showed no temporal trend, although it was the only dimension associated with seasonal climate: warmer previous summers and harvest-year autumns were associated with higher viability, and warmer harvest-year springs and summers with lower viability, and these associations were unchanged after controlling for annual crop size. Seed fill was not associated with seasonal climate, and seed mass showed at most a weak negative association with seasonal temperature that could not be assigned to any seasonal window. The decline in seed mass was not related to the masting history of the districts, nor to the local rate of decline in the interannual variability of seed production reconstructed from collection records. In cold storage, viability declined with storage duration and the rate of loss varied among harvest years more than the mean rate itself, but climate during seed development did not modify the rate of loss. Summer and autumn temperatures increased over the study period whereas spring temperature did not; the dimension of quality that responds to climate therefore remained stable, while the two dimensions that changed were at most weakly related to seasonal climate. It seems that the decline in seed mass and the increasing frequency of fill failure are driven by processes that seasonal climatic means do not capture, and their consequences for regeneration remain to be tested.

## Introduction

Forest regeneration depends on a long chain of processes, from flowering and pollination through seed development, dispersal and predation to germination, establishment and survival of seedlings. Seed production is the first link in this chain. Climate-driven changes in seed quantity, particularly the ongoing breakdown of masting in temperate trees (Ascoli et al., 2017; Bogdziewicz et al., 2020; Foest et al., 2025; Kelly and Sork, 2002; Pearse et al., 2016; Szymkowiak et al., 2024; Vacchiano et al., 2017), have received sustained research attention. The parallel question of whether climate also degrades the physiological quality of those seeds has been comparatively neglected (Harmel et al., 2026; Saatkamp et al., 2019), and seed counts alone tell us little about regeneration potential when the seeds being counted may be of progressively lower quality.

Pollination, fertilization, embryogenesis, and seed filling each unfold in different seasonal windows, and the same warming that triggers masting cues simultaneously alters the conditions experienced by developing seeds at every subsequent stage, with consequences that remain poorly resolved. Seed mass and seed viability, although both conventionally treated as components of seed quality, are determined by distinct physiological processes at different stages of seed development. Viability reflects the developmental integrity of the embryo, established during fertilization, embryogenesis, and maturation, and depends on cellular redox homeostasis (Bailly, 2004; Pukacka and Ratajczak, 2007; Ratajczak et al., 2019). Seed mass, in contrast, is set primarily during the later seed-filling period, when carbohydrate, lipid, and protein reserves are deposited, and is the principal determinant of initial seedling vigor (Moles and Westoby, 2006; Westoby et al., 1992). A third dimension, the proportion of filled seeds in a lot, is decided earlier still, at pollination and fertilization, and depends on pollen availability and on the loss of developing seeds to pre-dispersal predators such as *Cydia fagiglandana.* In masting species, pollination efficiency and predator satiation are density-dependent, so that the proportion of filled seeds is expected to vary with crop size through pollen coupling and predator satiation (Bogdziewicz et al., 2020; Pesendorfer et al., 2024). Because the three dimensions are built at different stages and through different processes, they should not be expected to share their climatic sensitivities, yet whether they actually diverge in response to climate has rarely been examined at scales adequate to resolve it.

A further consequence of changing seed quality concerns long-term storage, on which gene banks and forestry seed reserves rely (Kijowska-Oberc et al., 2021; Walters et al., 2005). Seeds matured under suboptimal conditions lose viability more rapidly in storage (Hay and Probert, 2013), so climate-driven change of initial quality may also change effective storage life, a question yet to be systematically tested using long-term banking datasets.

European beech (*Fagus sylvatica* L.), a foundational and economically important species of temperate European forests (Durrant et al., 2016; Ellison et al., 2005), provides a particularly suitable system in which to address these questions. It exhibits pronounced mast seeding governed by climatic cues, with a multi-year reproductive cycle whose successive stages can be aligned with seasonal climatic windows (Nussbaumer et al., 2016). Its seeds display intermediate storage behavior with limited longevity even under optimal cold storage (Pukacka and Ratajczak, 2007), and intensive management in Central Europe has generated long-term operational datasets. In Poland the species spans its natural range and reaches its northeastern margin (Jastrzębowski and Tarasiuk, 2010). A decline in the interannual variability and synchrony of seed production, termed masting breakdown, has been reported across Europe (Foest et al., 2024), including Poland (Foest et al., 2025).

The mechanisms that link seasonal climate to the three dimensions can be aligned with the reproductive calendar of beech. Warm and dry summers in the year preceding seed fall trigger flower initiation (Bogdziewicz et al., 2020; Vacchiano et al., 2017), and the same signal governs the allocation of carbohydrates to reproduction in the following year (Hacket-Pain et al., 2025). Warm previous summers may therefore act on seed quality in two directions: through the developmental quality of the flower buds initiated, which would favor viability, and through the depletion of stored reserves when abundant flowering follows, which would reduce the resources available per seed and hence seed mass (Kelly and Sork, 2002; Pearse et al., 2016). Spring of the harvest year encompasses pollination, fertilization and early embryogenesis; warm springs may shift pollen release relative to stigma receptivity (Kasprzyk et al., 2014; Simoleit et al., 2016), and thermal stress at this stage may disturb the redox homeostasis of the developing embryo, with irreversible oxidative damage to meristematic tissue (Kalemba et al., 2019; Pukacka and Ratajczak, 2007; Ratajczak et al., 2019). Summer and autumn of the harvest year cover seed filling and maturation, when seed mass is set and desiccation tolerance is acquired. Finally, because the proportion of filled seeds varies with crop size independently of climate, any climatic association at the year level may be confounded with crop size unless the latter is measured.

While studies on masting frequency continue to advance, laboratory assessments of individual-seed quality (germination tests, tetrazolium staining, excised embryo assays; ISTA Rules (International Seed Testing Association, 2026)) are rarely integrated with the large-scale, multi-year datasets that characterize contemporary masting research. Long-term operational seed monitoring conducted in Poland by the State Forests National Forest Holding (Państwowe Gospodarstwo Leśne Lasy Państwowe) has, however, generated a beech seed dataset of just such scale, spanning two decades and the full national range of the species, which we leverage here to test these questions.

Building on this dataset, we asked whether seasonal climate conditions acting at successive stages of the reproductive cycle are differentially associated with the three quality components, whether these dimensions changed over two decades, and whether the climate under which seeds developed is related to their viability during long-term storage. On the basis of the mechanisms outlined above, we hypothesized that: (I) warm summers in the year preceding seed fall would be positively associated with the viability of filled seeds and negatively, or not at all, with seed mass; (II) warm springs in the harvest year would be negatively associated with viability; (III) seed mass, seed fill and viability would be associated with non-overlapping seasonal windows, so that no single climatic driver would account for their temporal trajectories; and (IV) seeds developing under climatically unfavorable conditions would show lower viability throughout storage. Because crop size is known to influence the proportion of filled seeds (Pesendorfer et al., 2024), we tested the climatic associations with and without an independent measure of annual crop size.

## Material and Methods

### Study area

The study covers the entire range of European beech in Poland, spanning the species’ natural distribution in southern, western, and central regions of the country, as well as managed plantations established beyond the native range in northern Poland (Jastrzębowski and Tarasiuk, 2010). The dataset comprises 13,387 seed lots originating from 380 forest districts (Fig. 1), distributed from lowland to montane forests across an elevational gradient from near sea level to approximately 950 m a.s.l. The study area encompasses a broad climatic gradient typical of the Central European temperate zone, with mean annual temperature ranging from approximately 6 °C in the north-eastern lowlands to over 9 °C in the south-western and western parts of the country, and mean annual precipitation from around 500 mm in central Poland to over 1300 mm in the southern mountains (Kondracki, 2002; Woś, 2010).

**Figure 1.**
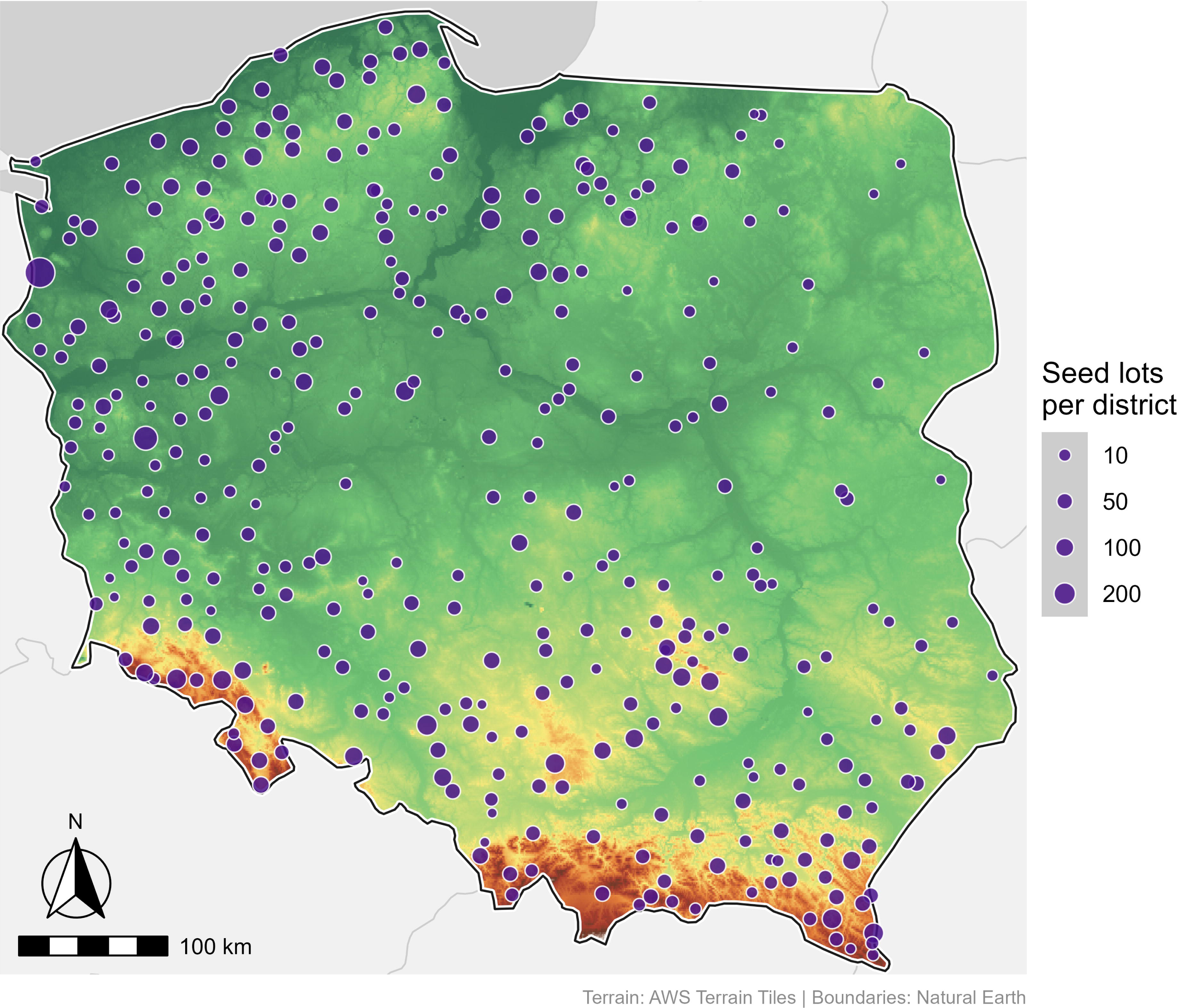
Distribution of European beech (*Fagus sylvatica* L.) seed lots across Poland, 1996 to 2024. Circles indicate forest districts contributing seed lots to the dataset (n = 13,387 from 380 forest districts), with circle size proportional to the number of seed lots collected from each district. The background shows topographic relief, with green colors indicating lowlands and orange-brown colors indicating highland and mountain areas. Temporal trends were analyzed for freshly harvested seed lots (storage duration = 0 years) with complete records, from 2004 to 2023 (n = 5,381 lots from 352 districts). Climate models were fitted to the subset with a crop index, from 2004 to 2020 (n = 4,698 lots from 312 districts). The storage analysis used 5,017 measurements of stored lots from 294 districts. Boundaries from Natural Earth; terrain from AWS Terrain Tiles.

### Data collection

#### Seed material

Seed material of European beech was collected between 1996 and 2024 by forest districts operating within the State Forests National Forest Holding. Each seed lot represents seeds collected from a single forest district during one harvest season, following the standard operational practice of bulk collection from multiple parent trees within the district. Seed lots were assessed at the Seed Assessment Stations of the State Forests National Forest Holding, which reported their results to the Forest Research Institute (Instytut Badawczy Leśnictwa, IBL). The method by which viability was assessed (staining of embryos, the germination test or the cutting test; see Seed quality assessment) is recorded for each lot and is referred to below as the testing method; 284 of 353 districts (80%) had lots tested with more than one method, so that the method effect is identifiable within districts, but in 2013 to 2015 all freshly harvested lots (n = 761) were assessed with the germination test, which was hardly used from 2016 onwards (6 of 2,510 lots; Table S1; see Limitations). The dataset includes both freshly harvested seeds (storage duration = 0 years) and seeds maintained in cold storage for up to 15 years. Beech seeds were dried to a moisture content of 9 to 11% in all facilities and stored at sub-zero temperatures following the protocol of Suszka et al. (2000). Stored lots are retested systematically and the lots that lose viability are successively withdrawn from the deposits, so that the lots tested after long storage are a selected subset of those originally stored. Because seed lots carry no individual identifier in the database, a stored lot cannot be linked to the fresh lot from which it originated; the storage analysis therefore compares stored lots with the mean of freshly harvested lots from the same district and harvest year (see below). Unless otherwise specified, the climate analyses described below are based on freshly harvested seed lots collected between 2004 and 2023 (n = 5,381 from 352 forest districts), the period for which complete E-OBS climate data were available.

#### Seed quality assessment

Purity was determined on a working sample of 100 g as the share by mass of pure seeds, that is, of well-developed, filled and undamaged seeds. Empty seeds and seeds damaged mechanically or by insects, including pre-dispersal predators such as *C. fagiglandana*, were excluded from the pure fraction. Viability was then evaluated on the pure fraction with one of three methods. In the tetrazolium test, the pericarp was removed and the seed coat was removed after soaking in water for 18 h. The embryos were then incubated for 10 to 12 h in the dark at 25 to 30 °C. The staining solution was 1% 2,3,5-triphenyltetrazolium chloride in water or in phosphate buffer at pH 6.5 to 7.5. Embryos stained completely, or with at least two thirds of the basal part of the cotyledons stained, were scored as viable. The germination test was carried out at 3 to 5 °C on four replicates of 100 pure seeds in a moist medium. Because of dormancy, the test may last up to about 150 days in beech (Suszka et al., 2000). In the cutting test, the proportion of visibly healthy, filled seeds was recorded. The three quantities were analyzed as separate dimensions of seed quality. Purity expresses the proportion by mass of filled seeds in the harvested material. Seed mass was recorded as the mass of 1000 seeds. Viability expresses the proportion of viable seeds among filled seeds and reflects embryo integrity. Their product, the proportion of viable seeds in the harvested material, is reported separately as an operational metric. Because the beta distribution is defined on the open interval (0,1), proportions were transformed before beta regression following Smithson and Verkuilen (2006): y′ = (y(n − 1) + 0.5)/n. This transformation affected 10 of 13,387 viability values (0.07%) and 443 purity values (3.3%) lying exactly at 0 or 1.

#### Climate data

Climatic data were obtained from the E-OBS gridded observational dataset (version 30.0e, ensemble mean) at a spatial resolution of 0.1° (Cornes et al., 2018). Climate variables were extracted for each forest district at its geographic centroid. The centroid of one coastal district (Międzyzdroje, four lots) fell on a grid cell dominated by sea and this district was excluded from climate analyses. We computed seasonal averages of mean temperature and sums of precipitation for both the harvest year and the year preceding seed collection, divided into spring (March–May), summer (June–August), and autumn (September–November). Terrain and country boundary data used for spatial visualizations were obtained from AWS Terrain Tiles via the elevatr R package (Hollister et al., 2023) and Natural Earth (naturalearthdata.com), respectively.

### Data quality control

The raw database contained 13,408 records. The analysis was restricted to seed lots collected in forest districts within Poland; 16 records from two suppliers outside Poland (one in Germany and one in the Czech Republic) were excluded on the basis of their administrative affiliation, together with three records without an identifiable forest district. In 43 records, values of purity, mass or viability had been converted into dates by spreadsheet autoformatting (e.g. 29.4 stored as 29 April); these values were recovered by reversing the conversion and were verified against the independently recorded number of seeds per kilogram. Two records tested with an X-ray method, too few to estimate a method effect, were removed. After applying these filters, the full dataset comprised 13,387 seed lots from 380 forest districts collected between 1996 and 2024. Climate analyses were further restricted to freshly harvested seed lots (storage duration = 0 years) collected between 2004 and 2023, the period for which complete E-OBS climate data were available, yielding 5,381 seed lots from 352 forest districts. The number of lots per year ranged from 3 to 822 (Table S1); because seed lots exist only where seeds were collected, years of low seed production are underrepresented.

### Data analysis

All analyses were performed in R version 4.5.3 (R Core Team, 2025). Linear mixed-effects models (LMM) were fitted using the lme4 and lmerTest packages (Bates et al., 2015; Kuznetsova et al., 2017), and generalized linear mixed-effects models (GLMM) with a beta distribution for proportions, or a binomial distribution for threshold exceedance, using glmmTMB (Brooks et al., 2017). All climatic predictors were standardized (mean subtracted and divided by the standard deviation, computed within the analytical dataset). Forest district and harvest year were included as random intercepts in all models. Testing method was included as a fixed effect in all models of viability; it was omitted from models of seed mass and purity, which are not determined by the viability test, after confirming that it had no effect on these responses (p ≥ 0.29) and did not change their temporal trends. Variance inflation factors were inspected for all multi-predictor models; because seasonal temperatures are correlated (r = 0.70 between previous-summer and spring temperature), VIF reached 5.2 to 6.2 for the three temperature terms of the reported viability model (2.3 to 2.5 after adjustment for degrees of freedom) and up to 7.4 in the full model, and climatic estimates are interpreted as conditional on the other seasonal terms. Model diagnostics used simulated scaled residuals in the DHARMa 0.4.7 package (Hartig, 2022), with tests of dispersion, uniformity (Kolmogorov-Smirnov D), outliers (bootstrap version) and spatial autocorrelation of district-level residuals (Moran’s I). Adjusted means and marginal predictions were computed with the emmeans 1.11.2.8 (Lenth et al., 2024) and ggeffects (Lüdecke, 2018) packages.

### Temporal trends in seed quality and climate

Seed lots collected in the same year are not independent observations of a temporal trend: the intraclass correlation of harvest year was 0.127 for seed mass, 0.141 for viability and 0.026 for purity, and omitting the year random effect from a trend model of seed mass underestimated the standard error of the linear slope 4.7-fold (41.7 vs 195.8 g). Temporal trends were therefore estimated with harvest year as the unit of replication. For each response, adjusted annual means were obtained from a mixed model with year as a factor and forest district as a random intercept (with testing method as a fixed effect for viability), and a linear trend was fitted to the 20 annual means by random-effects meta-regression (restricted maximum likelihood, metafor package; Viechtbauer, 2010), in which each annual mean was weighted by the inverse of the sum of its squared standard error and the among-year variance not explained by the trend. In this way years with many lots received more weight than years with few lots, but no single year could dominate the fit. Weighted regression with weights inversely proportional to the squared standard error alone, unweighted regression, restriction to years with at least 20, 50 or 100 lots and the omission of 2019, the year contributing 15% of the lots, were used as sensitivity analyses (Table S2). Because the distribution of purity is strongly left-skewed (median 0.963, first percentile 0.688) and the beta model of purity was overdispersed (dispersion ratio 1.33), the temporal change in seed fill was additionally described as the frequency of lots with fewer than 85% filled seeds, modelled with a binomial GLMM with year as a continuous fixed effect and district and year as random intercepts. Thresholds of 80% and 90% were used as sensitivity analyses. To describe changes in the whole distribution, the 10th, 50th and 90th percentiles of seed mass, the 5th and 10th percentiles of purity and the frequency of lots with fewer than 85% filled seeds were computed for each year with at least 20 lots (17 years) and regressed on year, each year carrying equal weight. Non-linear trajectories were explored with generalized additive models (GAMs) implemented with the mgcv package (Wood, 2017), with year fitted as a thin-plate regression spline and forest district as a random effect; the basis dimension was varied (k = 5 to 19) to check whether structure at the period of masting cycles was being smoothed away. Breakpoints were estimated on the annual means with the segmented package (Muggeo, 2008), compared with the linear model by AIC and Davies’ test, and their uncertainty was assessed by bootstrap (2000 resamples). Temporal trends in seasonal climate were tested by linear regression of national annual means on year, computed on a fixed grid of 379 forest districts × 20 years (7,580 district-years; one district had no climate data) and not on the districts and years in which seeds were collected, because the composition of the seed sample (3 to 219 districts per year) depends on crop size and would itself generate apparent climatic trends. Trends were also computed for each district separately (Fig. 2).

**Figure 2.**
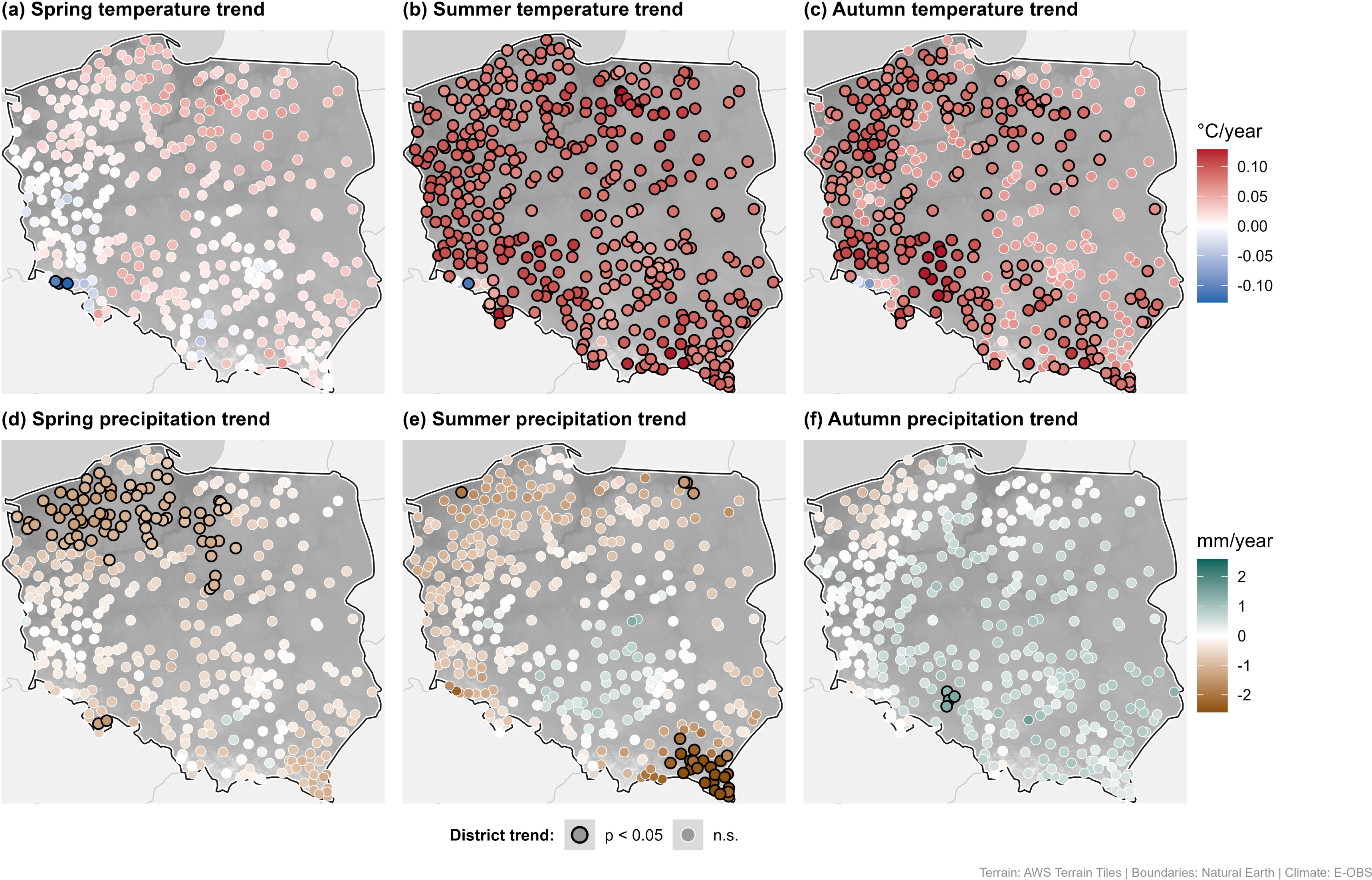
Trends in seasonal climate across Poland, 2004 to 2023. (a to c) Trends in mean temperature in spring (March to May), summer (June to August) and autumn (September to November); (d to f) trends in precipitation sums in the same seasons. Each circle represents one of 379 forest districts with a complete climate series. Colour indicates the change per year (°C year⁻¹ or mm year⁻¹) estimated by linear regression of the seasonal value on year, with red indicating warming and blue cooling in panels a to c, and green indicating wetting and brown drying in panels d to f. Circles with a black outline indicate trends significant at p < 0.05, and circles with a white outline non-significant trends. Background shading shows topographic relief in greyscale, with darker tones indicating lower elevations. The color scale is truncated at the 99th percentile of absolute trends; more extreme values are shown in the end colors. Climate data from the E-OBS gridded dataset (Cornes et al., 2018) at [0.1° × 0.1°] resolution.

### Climatic drivers of seed mass and viability

We modelled seed mass (LMM), purity (beta GLMM) and the viability of filled seeds (beta GLMM) as functions of seasonal climate. The twelve candidate predictors were seasonal means of temperature and sums of precipitation for spring, summer and autumn in the harvest year and in the preceding year. Instead of automated model selection, we compared a small set of a priori models derived from the reproductive phenology of beech: H1, previous-summer temperature and precipitation (flower initiation); H2, harvest-year spring temperature and precipitation (pollination and early embryogenesis); H4, harvest-year summer and autumn temperature and precipitation (seed filling and maturation); H124, the union of these three windows (eight predictors); a full model with all twelve predictors; and a null model with random effects only. Models were compared by AICc, and the H124 model is reported for all three responses with estimates and 95% confidence intervals for all predictors, including non-significant ones (Tables 1 and 2, Fig. 5); the full models are given in Table S3. Marginal and conditional R^2^ were computed following Nakagawa and Schielzeth (2013). To test whether the three dimensions respond to different climatic windows (hypothesis III) directly, and not by comparing the significance of predictors between separate models, we compared the standardized climatic coefficients between responses on a common scale (responses and predictors centered within district) and obtained confidence intervals for the pairwise differences by cluster bootstrap over forest districts (1000 resamples). Because the bootstrap over districts does not account for disturbances shared by all districts within a year, the procedure was repeated with harvest years as the resampled clusters, and two-sided bootstrap p-values were adjusted for the twelve predictors within each pair of responses with the Holm method (Holm, 1979). These analyses were restricted to freshly harvested seed lots for which the crop index described below was available (n = 4,698 lots from 312 districts and 17 harvest years, 2004 to 2020), so that all candidate models were fitted to the same data.

**Table 1.**
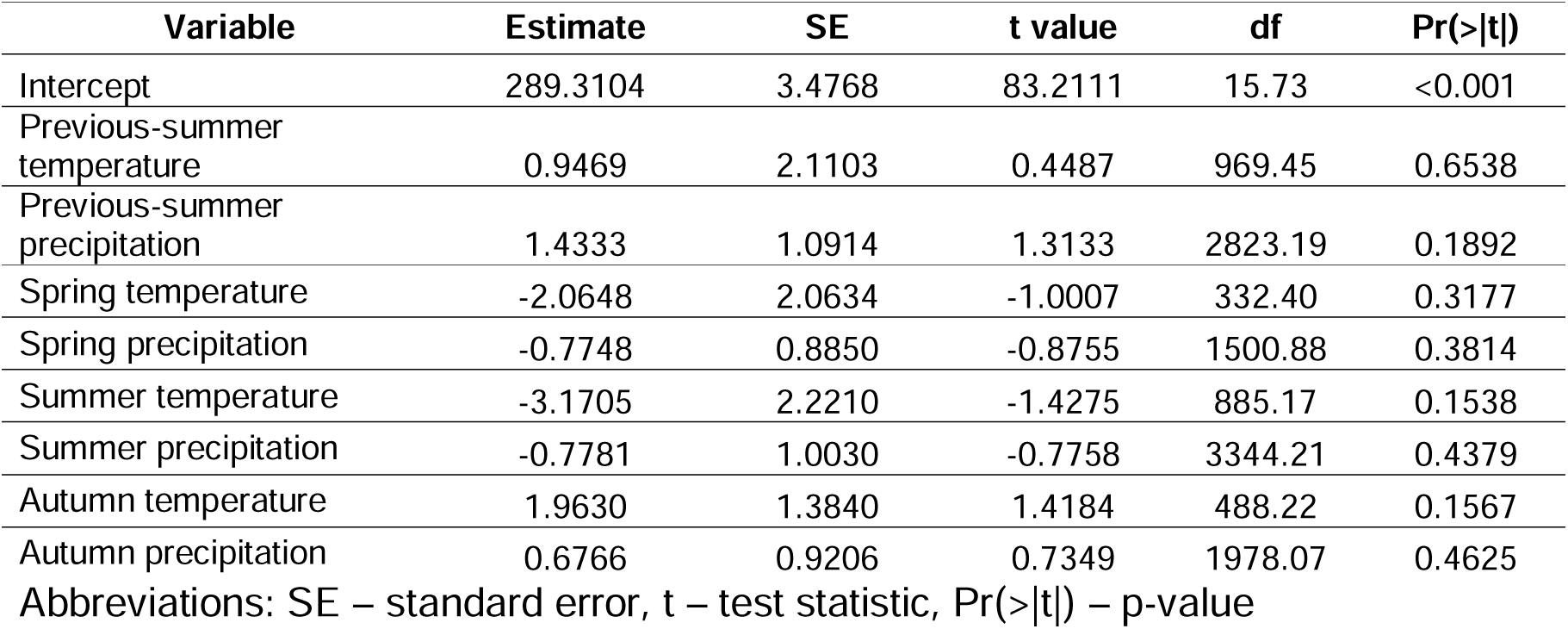
Linear mixed-effects model of seed mass (mass of 1000 seeds, g) as a function of seasonal climate during seed development (model H124). Predictors were standardized, so each estimate is the change in seed mass (g) per one standard deviation of the predictor. Forest district and harvest year were included as random intercepts, and the model was fitted by maximum likelihood to 4,698 freshly harvested seed lots from 312 forest districts and 17 harvest years (2004 to 2020). Degrees of freedom and p-values were obtained with the Satterthwaite approximation. Random effects SD: forest district = 5.16, harvest year = 13.1, residual = 39.6; R^2^m = 0.006, R^2^c = 0.118; AICc = 48,028.5, AICc of the null model (intercept and random effects only) = 48,028.0.

| Variable | Estimate | SE | t value | df | Pr(>\|t\|) |
| --- | --- | --- | --- | --- | --- |
| Intercept | 289.3104 | 3.4768 | 83.2111 | 15.73 | <0.001 |
| Previous-summer temperature | 0.9469 | 2.1103 | 0.4487 | 969.45 | 0.6538 |
| Previous-summer precipitation | 1.4333 | 1.0914 | 1.3133 | 2823.19 | 0.1892 |
| Spring temperature | -2.0648 | 2.0634 | -1.0007 | 332.40 | 0.3177 |
| Spring precipitation | -0.7748 | 0.8850 | -0.8755 | 1500.88 | 0.3814 |
| Summer temperature | -3.1705 | 2.2210 | -1.4275 | 885.17 | 0.1538 |
| Summer precipitation | -0.7781 | 1.0030 | -0.7758 | 3344.21 | 0.4379 |
| Autumn temperature | 1.9630 | 1.3840 | 1.4184 | 488.22 | 0.1567 |
| Autumn precipitation | 0.6766 | 0.9206 | 0.7349 | 1978.07 | 0.4625 |

Abbreviations: SE – standard error, t – test statistic, Pr(>|t|) – p-value

**Table 2.** Generalized linear mixed-effects model (beta distribution, logit link) of the viability of filled seeds as a function of seasonal climate during seed development (model H124). Predictors were standardized, so each estimate is the change in viability on the logit scale per one standard deviation of the predictor. The method of viability assessment was included as a fixed effect, with staining of embryos as the reference level. Forest district and harvest year were included as random intercepts, and the model was fitted to 4,698 freshly harvested seed lots from 312 forest districts and 17 harvest years (2004 to 2020). Random effects SD: forest district = 0.074, harvest year = 0.252; R^2^m = 0.164, R^2^c = 0.875; AICc = −10,434.2, AICc of the null model (intercept, method and random effects only) = −10,408.4.

| Variable | Estimate | SE | z | Pr(>\|z\|) |
| --- | --- | --- | --- | --- |
| Intercept | 1.6082 | 0.0679 | 23.6766 | <0.001 |
| Previous-year summer temperature | 0.1641 | 0.0327 | 5.0134 | <0.001 |
| Previous-year summer precipitation | 0.0156 | 0.0167 | 0.9353 | 0.3496 |
| Spring temperature | -0.1304 | 0.0332 | -3.9306 | <0.001 |
| Spring precipitation | 0.0042 | 0.0136 | 0.3083 | 0.7578 |
| Summer temperature | -0.1045 | 0.0341 | -3.0598 | 0.0022 |
| Summer precipitation | -0.0038 | 0.0154 | -0.2473 | 0.8046 |
| Autumn temperature | 0.0638 | 0.0218 | 2.9326 | 0.0034 |
| Autumn precipitation | 0.0054 | 0.0142 | 0.3821 | 0.7024 |
| Method: germination test (vs staining) | -0.1331 | 0.0450 | -2.9563 | 0.0031 |
| Method: cutting test (vs staining) | -0.0058 | 0.0283 | -0.2059 | 0.8369 |

Abbreviations: SE – standard error, z – test statistic, Pr(>|z|) – p-value

### Crop size and the scale of the climatic signal

To address the possibility that year-level climatic effects are confounded with annual crop size, we used five complementary approaches. First, annual seed collection data from the same operational source (Foest et al., 2025, 341 districts, 1988 to 2020) were matched to seed lots by district coordinates (325 districts matched), and a crop index was computed as the mass of seeds collected in a district and year relative to the demand set for that district, transformed as log(1 + x) and standardized within district. The index was available for 4,698 of the 5,381 freshly harvested lots. The climate models were fitted with and without this index, and the change in the climatic coefficients was used as a test of mediation by crop size. Second, each key climatic variable was decomposed into its national annual mean (between-year component) and the deviation of the district from that mean (within-year, spatial component), and both components were entered together in a model without the year random effect (Mundlak, 1978), to establish at which scale the climatic signal operates. Third, the viability model was refitted with year as a fixed effect, so that the climatic estimates rest exclusively on spatial contrasts within years. Fourth, the masting history of each district was derived from the same collection records for 1988 to 2020. A mast year was defined as a year in which the demand-corrected collection was positive and reached the upper quartile of the district series, and for each harvest year we computed the crop index of the preceding year, the number of years since the last mast year (truncated at 8 years) and the number of mast years in the five preceding years. These variables were added to the H124 models, the temporal trend in seed mass was refitted with and without them. Fifth, the number of freshly harvested lots per district and year (log-transformed and standardized within districts with at least five years of data) was used as an internal crop proxy, because it also covers 2021 to 2023, which are absent from the collection records; this proxy reflects collection effort as well as production (Spearman’s ρ = 0.41 with the crop index, n = 1,525 district-years). Finally, the H124 models were refitted to all 5,381 freshly harvested lots from 2004 to 2023, without the restriction to lots with a crop index.

### Masting variability and temporal trends in seed quality

To test directly whether the temporal trends in seed quality are related to masting breakdown, the interannual variability of seed production (CVp) was reconstructed for each district from the collection records following the procedure of Foest et al. (2025). CVp, the standard deviation divided by the mean of the mass of seeds collected, was computed in moving windows of 10 observations with a step of 5 years, and ln(CVp) was modelled with an LMM as a function of the last year of the window and the standardized mean seed demand in the window, with district as a random intercept. The estimated annual change in CVp was compared with that reported by Foest et al. (2025), and because successive windows overlap, the model was also refitted with a first-order autoregressive correlation structure (nlme package; Pinheiro and Bates, 2000). District-specific rates of change in CVp were obtained from a version of this model with a random slope for year, and their dependence on the reference temperature of the district was tested with a year × temperature interaction. As the reference temperature of a district, we used its mean temperature from March to November in 2004 to 2023, calculated from the same E-OBS data as the other climatic variables. The district-specific rates of change in CVp were then entered as a moderator of the temporal trend in each dimension of seed quality (year × rate interaction in the models described in Spatial structure of temporal trends); a positive interaction would indicate that seed quality declined faster where masting declined faster. The same test was carried out with the reference temperature and the rate of summer warming in 2004 to 2023 as moderators, and with an approximate change in CVp computed from the ratio of collection to demand in 1988 to 2004 and 2005 to 2020. As sensitivity analyses, the district rates of change in CVp were also estimated by least squares within each district, and the analysis was repeated without 2019. The p-values of all interaction terms were adjusted with the Holm method.

### Spatial structure of temporal trends

To assess whether the temporal trends were spatially structured, we fitted mixed models including year × latitude and year × longitude interactions as fixed effects, with random intercepts for forest district and harvest year and a random slope for year within district, so that the interactions are tested against the among-district variation in slopes (and with testing method as a fixed effect for viability). District-specific slopes were extracted from these models as conditional modes and summarized. For viability, the random slope for year within district could not be estimated (non-positive-definite Hessian), and the model was fitted with random intercepts for district and year only.

#### Relationships among the dimensions of seed quality

The relationships among seed mass, purity and the viability of filled seeds were assessed with Spearman’s rank correlation on the raw values of freshly harvested lots (n = 5,381), at the level of the individual lot and at the district-year level (means of lots), and viability was additionally modelled as a function of standardized seed mass in a beta GLMM with the random structure described above. Because multiplying two variables by a common factor induces correlation between them, the effect of correcting mass and viability by the common purity factor on their correlation was quantified separately (Table S4).

### Effects of storage on seed viability

To examine the loss of viability in storage and to test whether climate during seed development is related to it, we analyzed lots tested after 1 to 13 years of cold storage for which a fresh baseline was available (n = 5,017 measurements from 294 forest districts, forming 1,311 district × harvest-year cohorts). Most measurements were taken after short storage. The median storage duration was 2 years, 95.8% of the measurements were taken within 5 years, and 13 (0.3%) after more than 10 years. Because lots carry no identifier, the fresh baseline of a cohort was the mean viability of freshly harvested lots from the same district and harvest year, and the storage measurements are not repeated measures of the same lot. Viability of stored lots was modelled with a beta GLMM as a function of storage duration, the cohort baseline viability, the twelve seasonal climatic variables and testing method, with three biologically motivated storage duration × climate interactions (previous-summer temperature, and spring and summer temperature of the harvest year). The random structure comprised random intercepts for cohort, forest district and harvest year, with random slopes for storage duration at the cohort and harvest-year levels. The harvest-year random slope improved the fit substantially (ΔAIC = 8.5, likelihood ratio test p = 0.002) and its inclusion changed the inference on the interactions (see Results).

### Model diagnostics

Diagnostics of the reported models are given in Table S5. The beta model of purity was overdispersed, which motivated the binomial threshold model. Significant Kolmogorov-Smirnov tests reflected the discreteness of viability values, 66.1% of which were recorded as whole percentages.

### Software and visualization

All analyses, figures, and maps were produced in R version 4.5.3 (R Core Team, 2025) using the tidyverse, ggplot2, patchwork, sf, terra, tidyterra, emmeans, DHARMa and ggeffects packages. Elevation data were obtained via the elevatr package (Hollister et al., 2023) from AWS Terrain Tiles at approximately 250 m resolution. Country boundaries were sourced from Natural Earth (naturalearthdata.com).

## Results

### Temporal trends in seasonal climate

Over the period 2004 to 2023, summer and autumn warmed across Poland, whereas spring did not (Fig. 2, Fig. S1, Table S6). Mean summer temperature increased by 0.087 °C year⁻¹ (p = 0.0005) and autumn temperature by 0.074 °C year⁻¹ (p = 0.022). The trend in spring temperature was negligible (β = +0.016 °C year⁻¹, p = 0.67), and no seasonal precipitation variable showed a significant trend. The summer temperature trend was significant in 99% of districts and the autumn trend in 62%, against 1% for spring temperature (Fig. 2).

### Temporal trends in the three dimensions of seed quality

The three dimensions of seed quality changed over the 20-year period in different ways (Fig. 3). Adjusted annual mean seed mass declined by 1.48 g year⁻¹ (random-effects meta-regression on 20 annual means, p = 0.009). This corresponds to 29.6 g, or about 10% of the mean, over the study period (Fig. 3a, Table S2). The whole distribution shifted downwards, and the 10th, 50th and 90th percentiles of seed mass declined by 1.6 to 2.1 g year⁻¹ (p = 0.009 to 0.020) (Table S7). The decline remained significant in years with at least 20, 50 or 100 lots and after removing 2019, the year contributing 15% of the lots (β = −1.22 g year⁻¹, p = 0.014). It was not significant only in unweighted regression, which gives years with 3 to 13 lots the same weight as years with several hundred (β = −0.94 g year⁻¹, p = 0.16) (Table S2). No breakpoint could be located (Davies’ test p = 0.070; bootstrap 95% CI 2009 to 2019). The GAMs indicated year-to-year structure, but no slow trend hidden by smoothing (Fig. S2).

**Figure 3.**
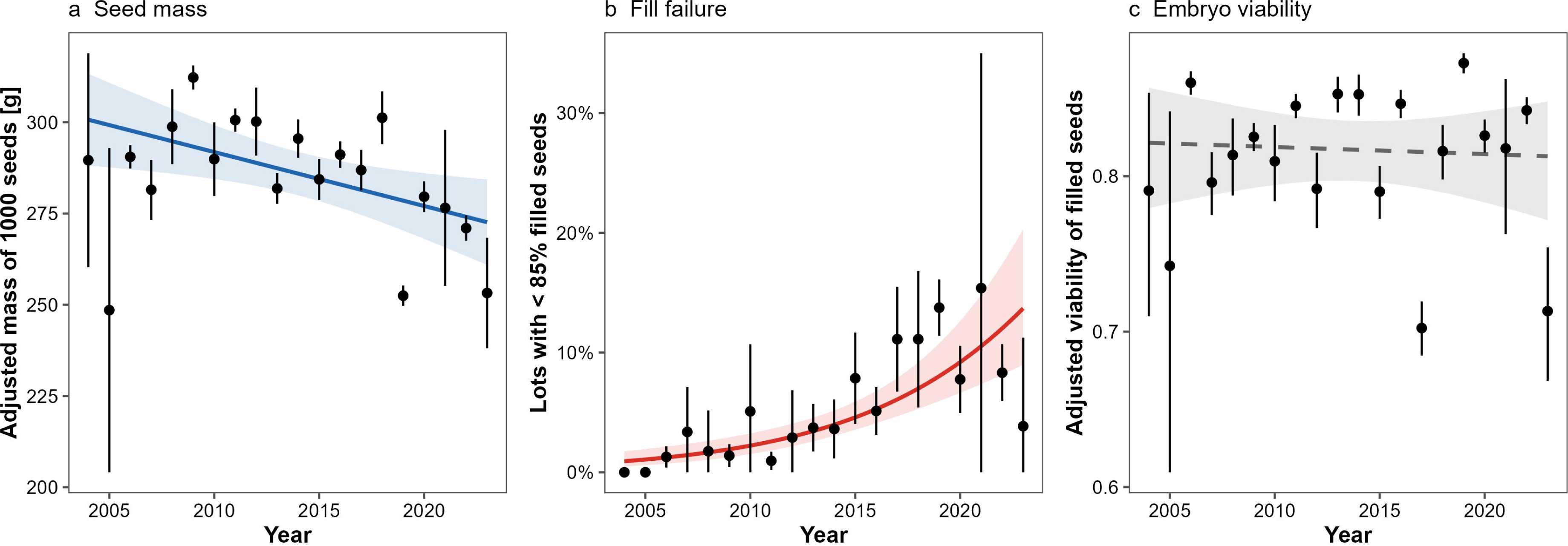
Temporal trends in three dimensions of seed quality of European beech (Fagus sylvatica L.) in Poland, 2004 to 2023. (a) Mass of 1000 seeds; (b) frequency of lots with fewer than 85% filled seeds; (c) viability of filled seeds. In panels a and c, points are annual means adjusted for forest district (mixed models with forest district as a random intercept) with 95% confidence intervals, and lines show the linear trend estimated by random-effects meta-regression on the 20 annual means, with 95% confidence bands. Viability was modelled on the logit scale and back-transformed for display. In panel b, points are the observed annual frequencies with 95% binomial confidence intervals, and the line shows the trend predicted by a binomial mixed model with forest district and harvest year as random intercepts, with its 95% confidence band. Solid lines indicate significant trends (seed mass, p = 0.009; fill failure, p < 0.001) and the dashed line a non-significant trend (viability, p = 0.80). Data from 5,381 freshly harvested seed lots from 352 forest districts.

The median proportion of filled seeds hardly changed (a decline of 0.5 percentage points over 20 years), and the decline in annual mean purity was not significant (β = −0.011 logit year⁻¹, p = 0.076). The lower tail of the distribution, however, widened (Fig. 4). The frequency of lots with fewer than 85% filled seeds increased from about 1.2% in 2006 to 13.7% in 2023 (binomial GLMM: β = 0.149 logit year⁻¹, p = 1.9 × 10⁻□; 95% CI for 2023, 9.0 to 20.3%; Fig. 3b). This trend did not depend on the threshold (80% or 90%) or on the inclusion of 2019.

**Figure 4.**
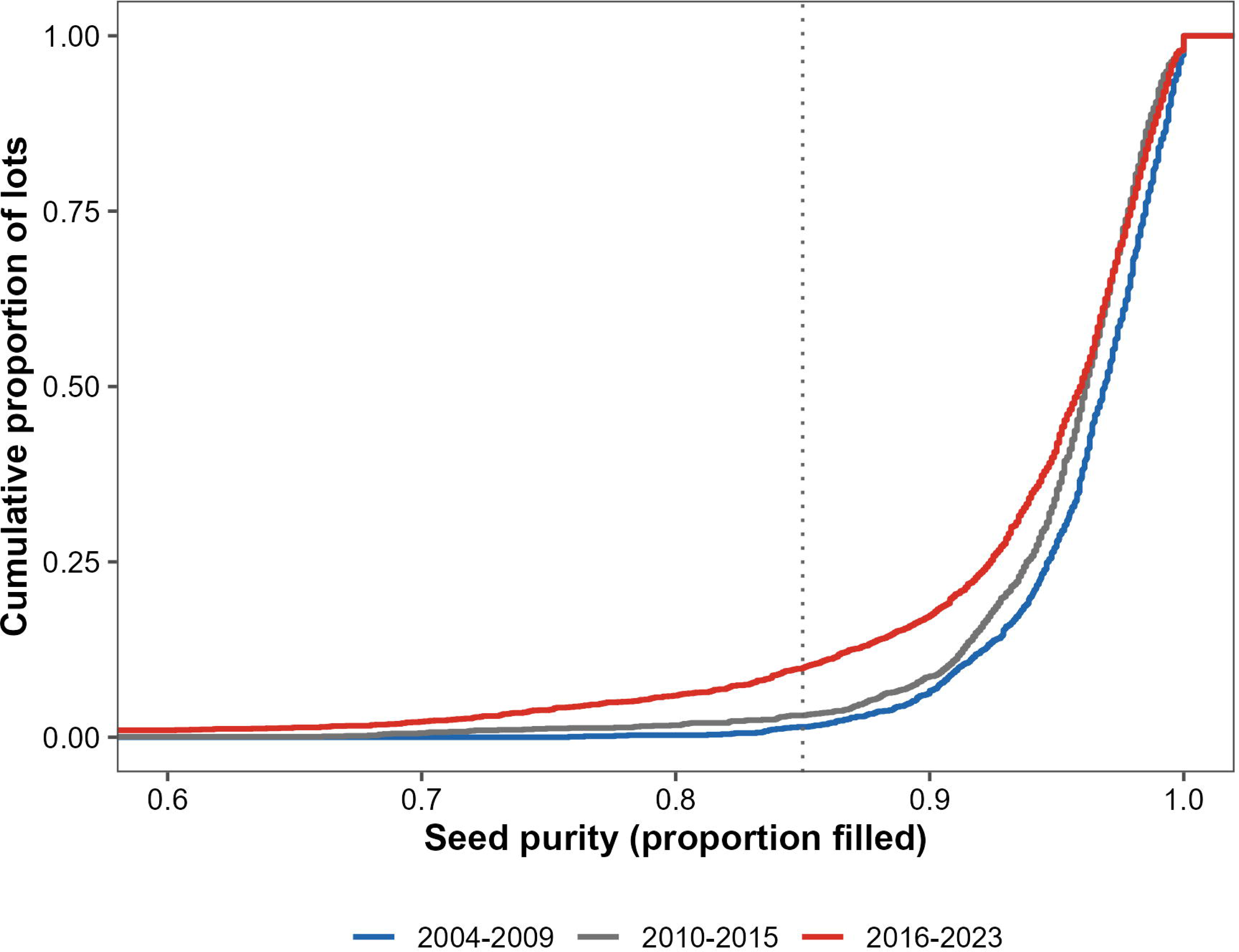
Distribution of seed purity (proportion of filled seeds) of European beech in three periods: 2004 to 2009 (1,354 lots), 2010 to 2015 (1,517 lots) and 2016 to 2023 (2,510 lots). Lines show the empirical cumulative distribution of purity across freshly harvested seed lots in each period. The dotted line marks the 85% threshold used to define lots with poor fill. The upper part of the distribution, including the median, hardly differed among periods, whereas the lower tail extended to lower values in the most recent period. Years contribute in proportion to their number of lots; the trend in poor fill was tested with a binomial mixed model with harvest year as a random effect (Fig. 3b). The x-axis is truncated at 0.6; lots with lower purity (0.5% of all lots) are included in the cumulative proportions but not displayed.

The viability of filled seeds showed no trend (β = −0.003 logit year⁻¹, p = 0.80; Fig. 3c). The same was true when only lots assessed by staining of embryos were analysed (β = −0.004, p = 0.77).

Seed mass and the viability of filled seeds were essentially uncorrelated at the level of lots (Spearman’s ρ = −0.056, n = 5,381). At the district-year level, all three pairwise correlations were weak (|ρ| ≤ 0.11, n = 2,082) (Table S8).

The decline in seed mass was spatially uniform. Neither the year × latitude nor the year × longitude interaction was significant for any dimension (p ≥ 0.05) (Table S9). The district-specific slopes of seed mass were negative in all 352 districts (median −1.49 g year⁻¹).

### Climate associations with seed mass, seed fill and viability

Among the a priori models, the H124 model was retained for viability (ΔAICc = 26 relative to the null model), whereas for purity the null model had the lowest AICc (Tables 1 and 2, Table S3, Fig. 5). For seed mass, each single-window model improved on the null model (ΔAICc = −4.7 to −5.4), with negative temperature coefficients of −2.5 to −4.0 g per standard deviation. Their union H124 did not improve on the null model, and none of its predictors differed from zero (Table 1). Because seasonal temperatures are correlated, the weak negative association of seed mass with temperature could not be assigned to any seasonal window.

**Figure 5.**
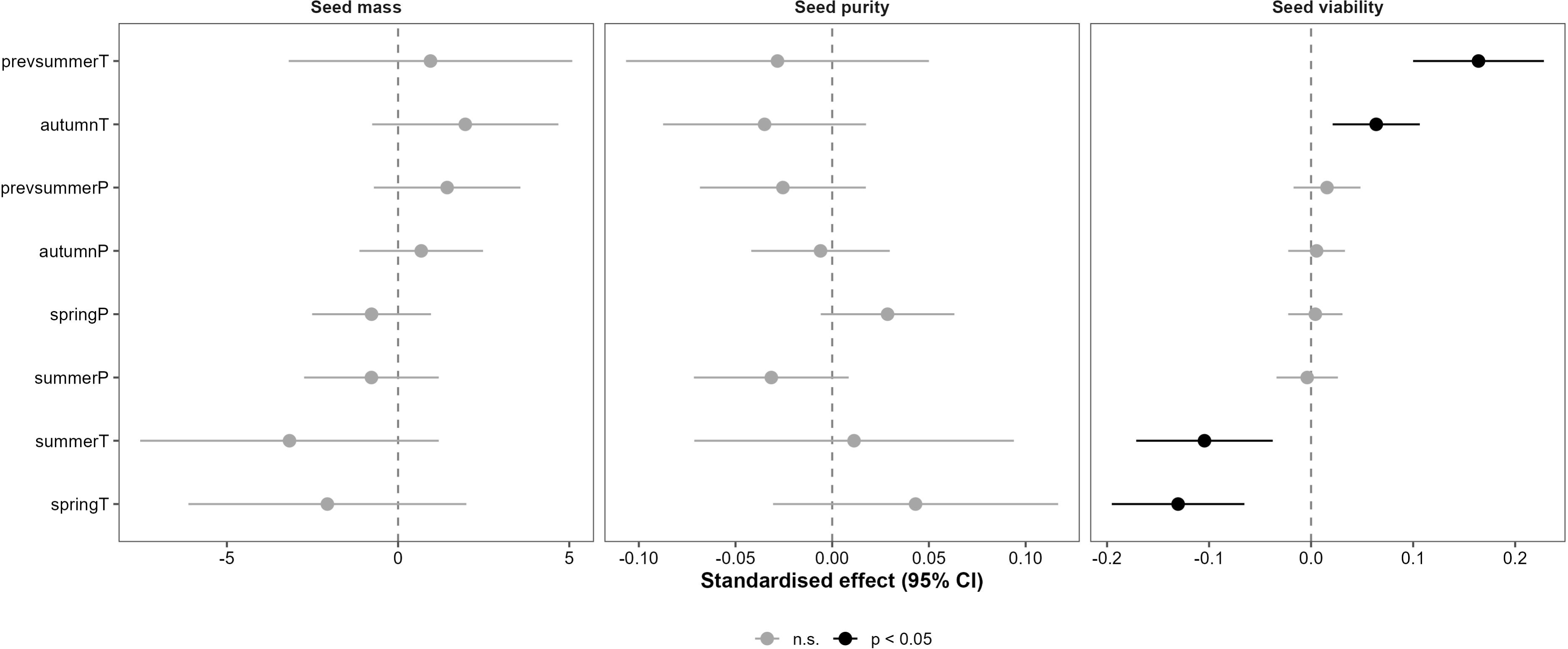
Associations of seasonal climate during seed development with seed mass, seed purity and the viability of filled seeds of European beech (models H124; Tables 1 and 2, Table S3). Points show the estimated effect of a one standard deviation increase in each predictor, and horizontal lines its 95% confidence interval. Black symbols indicate effects significant at p < 0.05, grey symbols non-significant effects. Effects are expressed in grams of 1000 seeds for seed mass and on the logit scale for purity and viability, so their magnitudes are comparable within, but not between, panels. Predictors: temperature (T) and precipitation (P) in the summer of the year preceding seed fall (prev) and in spring, summer and autumn of the harvest year. Models were fitted to 4,698 freshly harvested seed lots from 312 forest districts (2004 to 2020), with forest district and harvest year as random intercepts; the viability model also included the method of viability assessment.

The H124 model explained 16.4% of the variance in viability (R^2^m = 0.164, R^2^c = 0.875) and 0.6% of the variance in seed mass (R^2^m = 0.006). In the viability model, all four temperature terms were significant, and their signs alternated between seasons (Table 2). Previous-summer temperature (β = +0.164, 95% CI 0.100 to 0.228) and harvest-year autumn temperature (β = +0.064, CI 0.021 to 0.106) were positively associated with viability. Harvest-year spring temperature (β = −0.130, CI −0.195 to −0.065) and summer temperature (β = −0.104, CI −0.171 to −0.038) were negatively associated. No precipitation term differed from zero. Lots assessed with the germination test had lower viability than lots assessed by staining of embryos (β = −0.133, p = 0.003).

The effect of previous-summer temperature was conditional on spring temperature, with which it is correlated (r = 0.70); entered alone, it was zero (p = 0.999). The four temperature effects were stable in six sensitivity analyses, including year as a fixed effect and lots assessed by staining only (Table S10). With the bootstrap over districts, the climatic coefficients for viability differed from those for purity and from those for seed mass for seven of the twelve predictors each (Table S11). With the more conservative bootstrap over harvest years, two and four of these differences remained. Only the differences in previous-summer temperature and autumn precipitation were supported in both comparisons involving viability under all criteria.

### Crop size, masting history and masting variability

Adding the crop index to the viability model changed the climatic coefficients by at most 5%, and no coefficient changed sign or significance. The index itself was not associated with viability, seed mass or purity in the models with harvest year as a random effect (Table S12). For previous-summer and spring temperature, the climatic signal was present within years as well as between years (within-year β = +0.081 and −0.098, both p < 0.001) (Table S13). The climatic results were the same for all 5,381 lots and with an internal crop proxy covering 2004 to 2023 (Table S14).

Masting history was not associated with seed mass or purity (p ≥ 0.33). It included the crop of the preceding year, the time since the last mast year and the number of mast years in the five preceding years. Masting history did not change the temporal slope of seed mass (−0.97 and −0.96 g year⁻¹ without and with it) (Table S15). Viability was slightly lower after large crops in the preceding year (β = −0.031, 95% CI −0.057 to −0.004).

Our reconstruction reproduced the published decline in masting variability. CVp declined by 0.99% per year (95% CI 0.81 to 1.16%), against 1.06% reported by Foest et al. (2025), and the decline was weaker in warmer districts (Table S16). The temporal trends in seed quality did not depend on the rate of this decline (n = 5,234 lots from 311 districts). For seed mass, the year × rate interaction was β = −0.08 g year⁻¹ per standard deviation (95% CI −0.33 to 0.17, p = 0.54), which excludes a moderation larger than about 23% of the mean slope. The interactions for viability and for the frequency of poorly filled lots were not significant either (p ≥ 0.50). For mean purity, a nominally significant interaction in the opposite direction did not survive correction for multiple testing (Holm-adjusted p = 0.62). Neither the reference temperature nor the rate of summer warming of the districts modified any trend (p ≥ 0.63; Fig. S3, Table S16).

### Storage performance and climatic legacy effects

Viability declined with storage duration (β = −0.109 per standard deviation of storage duration, equal to 1.55 years, p = 0.015; Fig. 6, Table 3). Harvest years differed in the rate of loss more than the mean rate itself. The standard deviation of the harvest-year random slope for storage duration (0.128) exceeded the mean effect. In a model without this random slope, the interactions of storage duration with harvest-year spring temperature (β = +0.056, p = 0.001) and summer temperature (β = −0.047, p = 0.008) were significant. After allowing harvest years to differ in their rate of loss, none of the three interactions was supported (spring temperature β = +0.028, p = 0.32; summer temperature β = −0.009, p = 0.80; previous-summer temperature β = −0.029, p = 0.39). Climate during seed development was thus related to the level of viability in storage and not to the rate at which it was lost.

**Figure 6.**
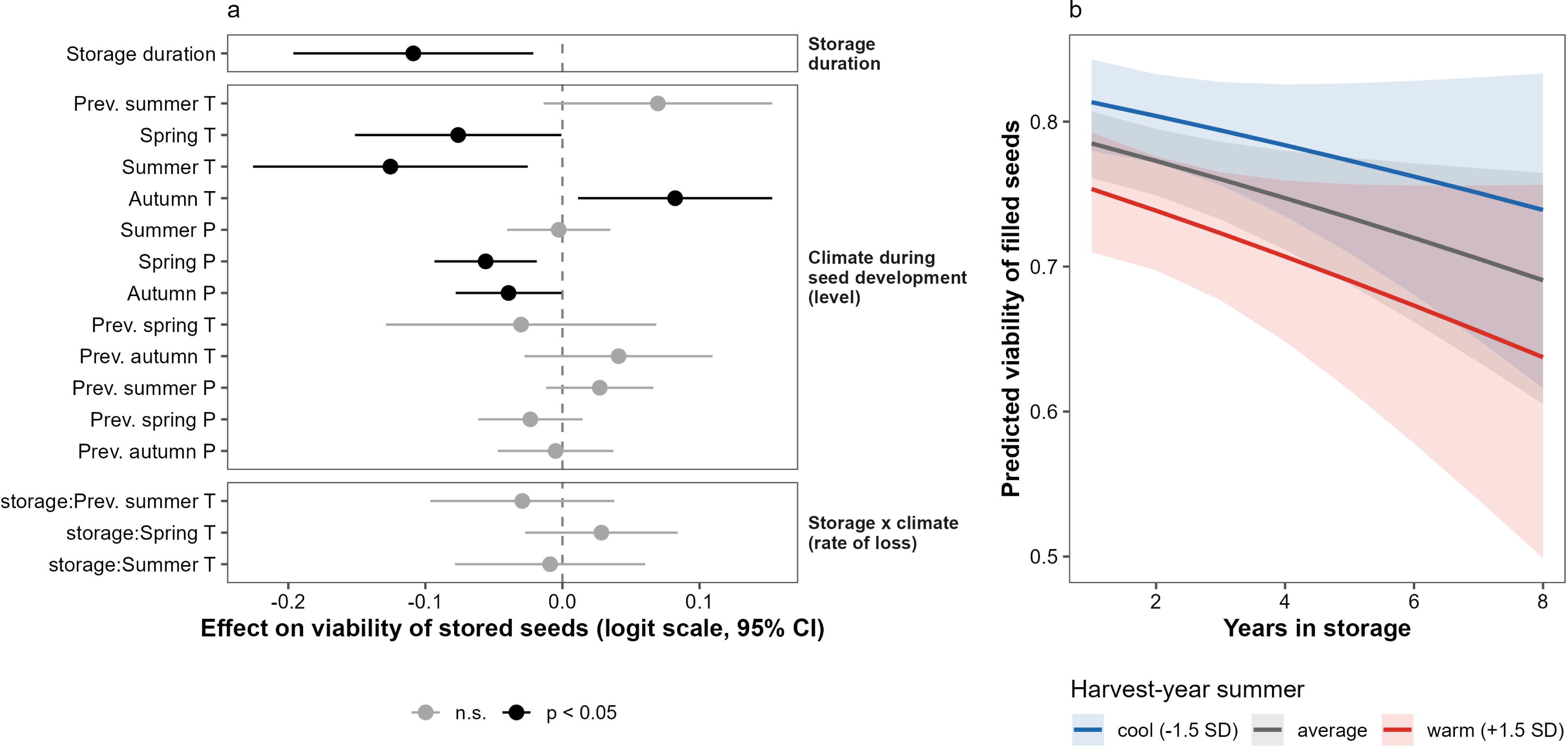
Viability of filled seeds in stored seed lots of European beech in relation to storage duration and to climate during seed development (Table 3). (a) Estimated effects with 95% confidence intervals on the logit scale, per one standard deviation of each predictor, grouped into storage duration, climate during seed development (effects on the level of viability) and interactions of storage duration with climate (effects on the rate of viability loss). Black symbols indicate effects significant at p < 0.05, grey symbols non-significant effects. T, temperature; P, precipitation; Prev., year preceding seed fall; other seasons refer to the harvest year. (b) Predicted viability during the first eight years of storage for harvest years with cool (−1.5 SD), average and warm (+1.5 SD) summers, with all other predictors at their means; shaded bands are 95% confidence intervals. Lines are nearly parallel on the logit scale; their divergence on the proportion scale results from the back-transformation and does not indicate a difference in the rate of loss. The model was fitted to 5,017 measurements of stored lots from 1,311 cohorts (forest district × harvest year) in 294 forest districts, with random intercepts and random slopes for storage duration at the cohort and harvest-year levels.

**Table 3.** Generalized linear mixed-effects model (beta distribution, logit link) of the viability of filled seeds in stored seed lots as a function of storage duration and seasonal climate during seed development. Predictors were standardized, so each estimate is the change in viability on the logit scale per one standard deviation of the predictor (one standard deviation of storage duration equals 1.55 years). Interactions of storage duration with climate test whether climate during seed development modified the rate of viability loss. The method of viability assessment was included as a fixed effect, with staining of embryos as the reference level. The cohort (forest district × harvest year) was included as a random intercept with a random slope for storage duration, forest district as a random intercept, and harvest year as a random intercept with a random slope for storage duration. The model was fitted to 5,017 measurements of stored lots from 1,311 cohorts, 294 forest districts and 19 harvest years. Random effects SD: cohort intercept = 0.135, cohort slope = 0.068, forest district = 0.042, harvest-year intercept = 0.248, harvest-year slope = 0.128; R^2^m = 0.178, R^2^c = 0.843; AICc = −8,048.9.

| Variable | Estimate | SE | z | Pr(>\|z\|) |
| --- | --- | --- | --- | --- |
| Intercept | 1.2520 | 0.0682 | 18.3614 | <0.001 |
| Storage duration | -0.1088 | 0.0446 | -2.4363 | 0.0148 |
| Previous-year summer temperature | 0.0696 | 0.0426 | 1.6358 | 0.1019 |
| Spring temperature | -0.0761 | 0.0384 | -1.9798 | 0.0477 |
| Summer temperature | -0.1255 | 0.0511 | -2.4537 | 0.0141 |
| Baseline viability of the cohort | -0.0043 | 0.0118 | -0.3648 | 0.7153 |
| Autumn temperature | 0.0822 | 0.0361 | 2.2762 | 0.0228 |
| Summer precipitation | -0.0027 | 0.0192 | -0.1400 | 0.8887 |
| Spring precipitation | -0.0560 | 0.0191 | -2.9403 | 0.0033 |
| Autumn precipitation | -0.0393 | 0.0197 | -1.9994 | 0.0456 |
| Previous-year spring temperature | -0.0302 | 0.0503 | -0.5994 | 0.5489 |
| Previous-year autumn temperature | 0.0410 | 0.0350 | 1.1722 | 0.2411 |
| Previous-year summer precipitation | 0.0272 | 0.0200 | 1.3645 | 0.1724 |
| Previous-year spring precipitation | -0.0234 | 0.0194 | -1.2056 | 0.2280 |
| Previous-year autumn precipitation | -0.0050 | 0.0216 | -0.2318 | 0.8167 |
| Method: germination test (vs staining) | -0.1159 | 0.0346 | -3.3523 | <0.001 |
| Method: cutting test (vs staining) | -0.0181 | 0.0280 | -0.6456 | 0.5185 |
| Storage duration × previous-year summer temperature | -0.0293 | 0.0342 | -0.8548 | 0.3927 |
| Storage duration × spring temperature | 0.0284 | 0.0284 | 1.0017 | 0.3165 |
| Storage duration × summer temperature | -0.0090 | 0.0354 | -0.2539 | 0.7996 |

Abbreviations: SE – standard error, z – test statistic, Pr(>|z|) – p-value

The most consistent associations were with harvest-year summer temperature (β = −0.125, CI −0.226 to −0.025) and spring precipitation (β = −0.056, CI −0.093 to −0.019). The fixed effects explained 17.8% of the variance in viability of stored lots (R^2^m = 0.178). The baseline viability of the cohort had no effect once the cohort random effect was included (p = 0.72).

## Discussion

Using a 20-year, nation-wide dataset, we followed three dimensions of beech seed quality separately and found that they changed over time and were related to climate in different ways. Hypotheses I and II were supported for viability: warmer previous summers were associated with higher, and warmer springs of the harvest year with lower viability of filled seeds. Hypothesis III was supported: the three dimensions were almost uncorrelated, and their climatic coefficients differed, most consistently for previous-summer temperature and autumn precipitation. Hypothesis IV was supported only in part, because climate during seed development was related to the level of viability in storage but not to the rate of its loss.

The two dimensions that changed, seed mass and seed fill, showed at most a weak climatic signal, whereas viability, the only dimension clearly associated with climate, did not change. Moreover, the seasons that warmed in Poland, summer and autumn, have associations of opposite sign with viability, and spring did not warm. We therefore do not attribute the trends in seed quality to climate change. Seed quality in beech is changing along axes that seed counts do not register, and seasonal climatic means do not explain why.

The positive association of previous-summer temperature with viability was conditional on spring temperature. Elevated temperatures in the prior summer trigger flower initiation in beech (Vacchiano et al., 2017), and Bogdziewicz et al. (2020) showed that warming of this cue disrupts the interannual variability of seed production. Our result suggests that the same cue may also be related to the developmental quality of the flower buds. This is consistent with the response of beech to the climatic conditions of the previous growing season reported for radial growth (Stojanović et al., 2018), but the mechanism remains a hypothesis. For seed mass, the same window had at most a weak negative effect, so the resource dilution mechanism of Kelly and Sork (2002) and Pearse et al. (2016) was barely detectable in our data.

Like Kondrat et al. (2026), we found no evidence that seed mass decreases with crop size. Their estimate, however, comes from individual trees sampled in two years, and ours from district-level crops over 17 years, with an index that reflects collection effort as well as production (Table S12). The positive association of the crop index with viability appeared only between years (Table S13). It would be consistent with pollen coupling (Pesendorfer et al., 2024), but it cannot be separated from other year-level factors. The slightly lower viability after large crops in the preceding year (Table S15) may reflect the depletion of reserves and should be regarded as a hypothesis.

Spring encompasses pollination, fertilization and early embryogenesis, and we regard two explanations of its negative association with viability as plausible. The first is a phenological mismatch between pollen release and stigma receptivity (Kasprzyk et al., 2014; Simoleit et al., 2016). The second is thermal stress disturbing the redox homeostasis of the embryo, consistent with the loss of viability caused by an imbalance of reactive oxygen species in the radicle meristem of beech embryos (Kalemba et al., 2019). Both explanations remain hypotheses that motivate experimental work on developing seeds.

The decline in seed mass coincided with the period in which Foest et al. (2025) reported masting breakdown in Poland, that is, a decline in the interannual variability and synchrony of seed production (Foest et al., 2024). We tested this coincidence directly and did not find a link. The resource depletion mechanism proposed by Hacket-Pain et al. (2025) requires shorter intervals between large crops or a greater cumulative allocation to reproduction, which masting breakdown does not necessarily imply. Yet seed mass did not decline faster where masting declined faster, and it was not related to masting history, that is, to the time since the last mast year or the number of mast years in the preceding five years. Neither masting breakdown nor the frequency of large crops therefore explains the decline in seed mass at the scale of forest districts, and its cause remains unknown.

The increase in poorly filled lots, with a nearly unchanged median fill, points to a different process. Fill failure arises from unsuccessful pollination and from pre-dispersal predation, and both are density-dependent in beech. Pesendorfer et al. (2024) showed that pollination efficiency and predator satiation fall steeply at low levels of reproduction, and low seed years, which starve seed predators, have become rarer (Foest et al., 2024). Seeds may thus be empty because pollination failed, or damaged by predators such as *C. fagiglandana*. The database records only the share of pure seeds, so we cannot tell which of the two causes underlies the increase.

Whether lighter seeds and more frequent fill failure affect regeneration cannot be decided from our data. Pawłowski et al. (2024) showed that smaller beech seeds germinate later and produce smaller seedlings, and Graf et al. (2026) reported that seed size and the proportion of fertile seeds promote establishment at range edges. A weaker start at the seed stage may also be compensated later in development. Beech shows high phenotypic plasticity, and Chmura et al. (2024) found that the growth and productivity of beech populations respond plastically to climatic transfer at the north-eastern border of the range. Therefore, it remains unclear whether lighter seeds leave a lasting mark on the next generation of trees.

Viability declined with storage duration, and the rate of loss differed among harvest years more than the mean rate itself (Table 3). Once this variation was allowed for, climate during seed development was related to the level of viability in storage and not to the rate of its loss. Seeds from years with warm summers and wet springs had lower viability throughout storage. This is consistent with, but does not demonstrate, the accumulation of oxidative damage during development (Ratajczak et al., 2019). Lots were tested during storage up to 15 years, whereas Pukacka and Ratajczak (2007) reported only 20% germination after 10 years of optimal storage. Such lots were rare (0.3% of measurements), and lots that lose viability are withdrawn from the deposits, so our estimates describe mainly the first years of storage. Even within these first years, however, seeds from different harvest years differed consistently in viability. Whether these differences are set already during seed development remains to be tested. Measurements of mitochondrial respiration in beech seeds (Fuchs et al., 2023) offer a way to test whether oxidative status at harvest may predict them.

Our dataset originates from operational monitoring, and several constraints temper causal inference. Seed lots exist only where seeds were collected, so years of low seed production are underrepresented and the data describe seed quality in collected crops. In 2013 to 2015 all fresh lots were assessed with the germination test, so method and period are partly aliased, although the results were unchanged for a single method (Table S10). Climate was assigned to district centroids, which would bias climatic effects towards zero. The alternating signs of consecutive seasonal terms suggest that daily extremes, such as late frost or heat during seed filling, may be more relevant than seasonal means. The crop index, the masting history and the rates of masting decline were derived from collection records, which reflect collection effort as well as production. Finally, the physiological mechanisms discussed above remain hypotheses.

Our findings show that laboratory measurements of seed quality can be integrated with landscape-scale monitoring. In Polish beech, seed mass and the frequency of poorly filled lots have changed over two decades in ways that seed counts do not register. These changes were explained neither by seasonal climate nor by masting history. Whether they affect regeneration remains to be established, and this will require linking seed quality records to seed production, predation and establishment data at the same sites.

## Acknowledgements

We acknowledge the E-OBS dataset from the EU-FP6 project UERRA (http://www.uerra.eu) and the Copernicus Climate Change Service, and the data providers in the ECA&D project (https://www.ecad.eu). We thank the State Forests National Forest Holding (Państwowe Gospodarstwo Leśne Lasy Państwowe) for providing access to the long-term seed quality monitoring dataset.

## Author contributions: CrediT

**Hanna Fuchs:** Data curation, Formal analysis, Investigation, Methodology, Visualization, Project administration, Writing – original draft, Writing – review & editing. **Marcin K. Dyderski:** Data curation, Formal analysis, Investigation, Methodology, Visualization, Writing – original draft, Writing – review & editing. **Szymon Jastrz**ę**bowski:** Data curation, Investigation, Writing – review & editing. **Ewelina Ratajczak:** Conceptualization, Supervision, Data curation, Investigation, Writing – original draft, Writing – review & editing.

## Data availability statement

The data and R code supporting the findings of this study are openly available in Figshare at https://doi.org/10.6084/m9.figshare.32346645. The repository includes seed quality data, climatic variables, and analysis scripts with full specifications of the linear mixed-effects models and segmented regression analyses. The repository will be made publicly accessible upon publication of this article.

