## Supplementary material for "Beyond seed counts: what two decades of seed testing reveal about European beech in Poland": Supplementary Figures.docx

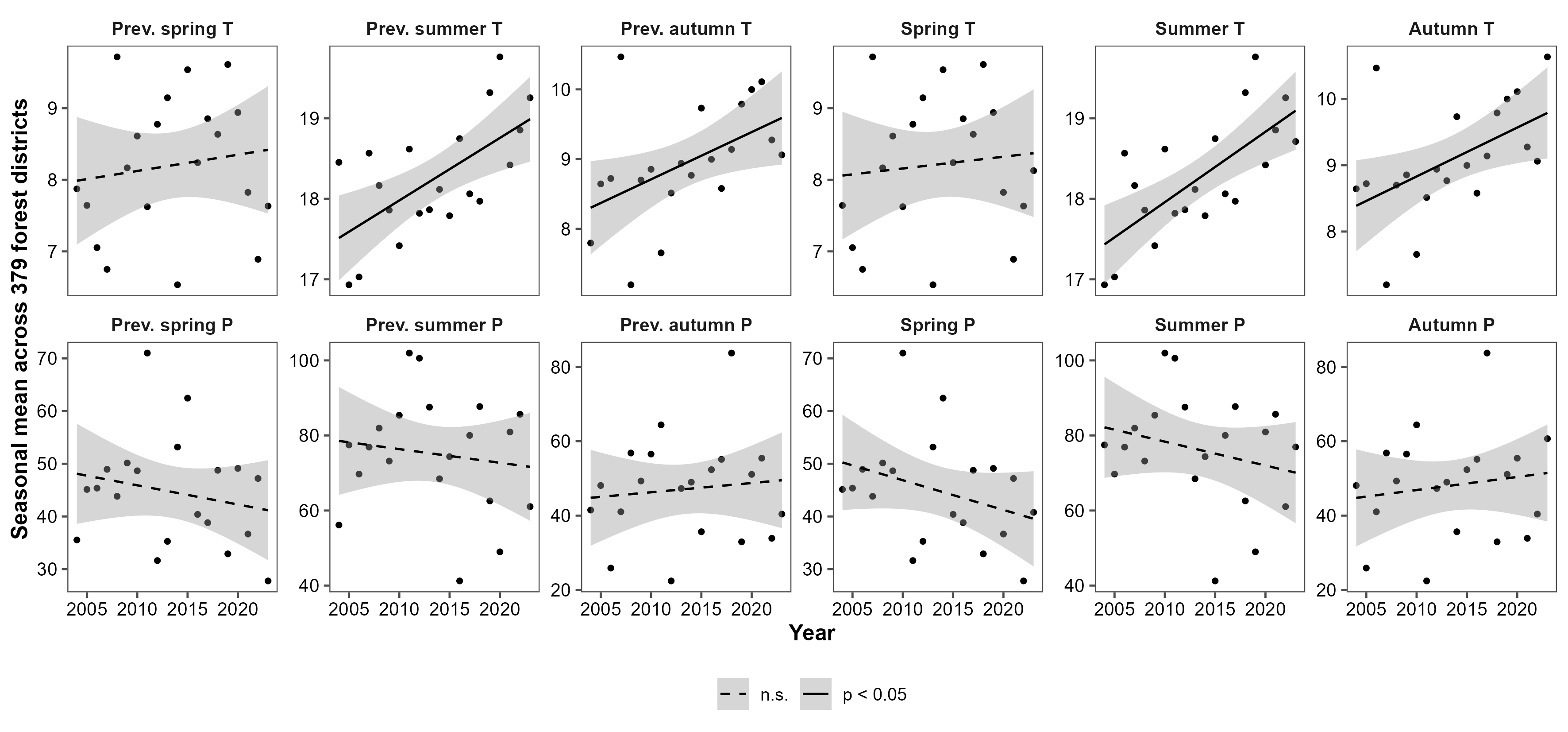

Fig. S1. National means of seasonal temperature and precipitation in the harvest year and in the preceding year, 2004 to 2023, calculated on a fixed grid of 379 forest districts. Lines show linear trends with 95% confidence bands; solid lines indicate trends significant at p < 0.05 and dashed lines non-significant trends (Table S6).

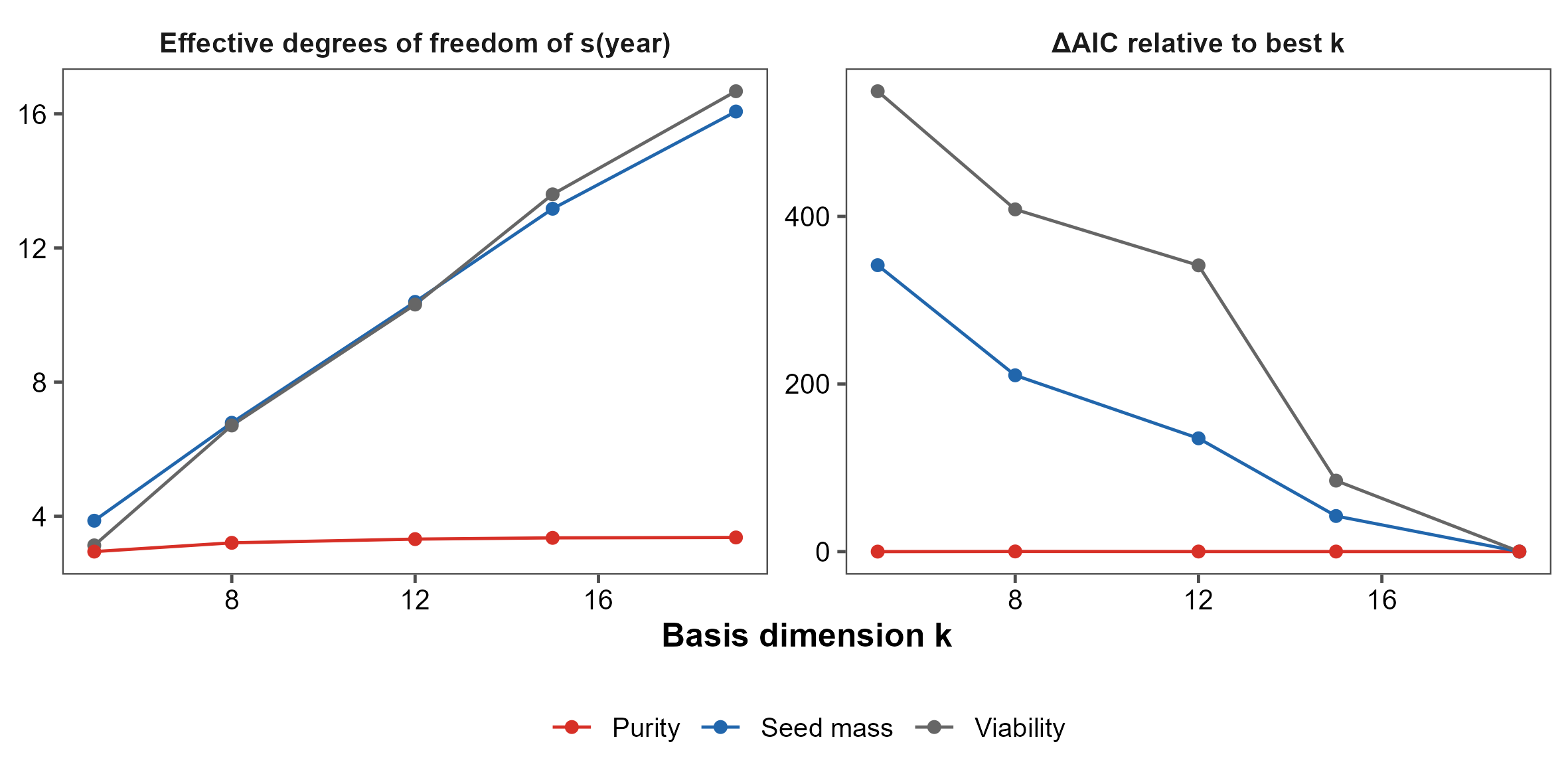

Fig. S2. Sensitivity of the generalized additive models of temporal trends in seed mass, viability and purity to the basis dimension (k) of the year smooth. Left: effective degrees of freedom of the year smooth; right: difference in AIC relative to the model with the lowest AIC. For seed mass and viability, the effective degrees of freedom increased with k and AIC decreased throughout, indicating year-to-year structure and no slow trend hidden by smoothing.

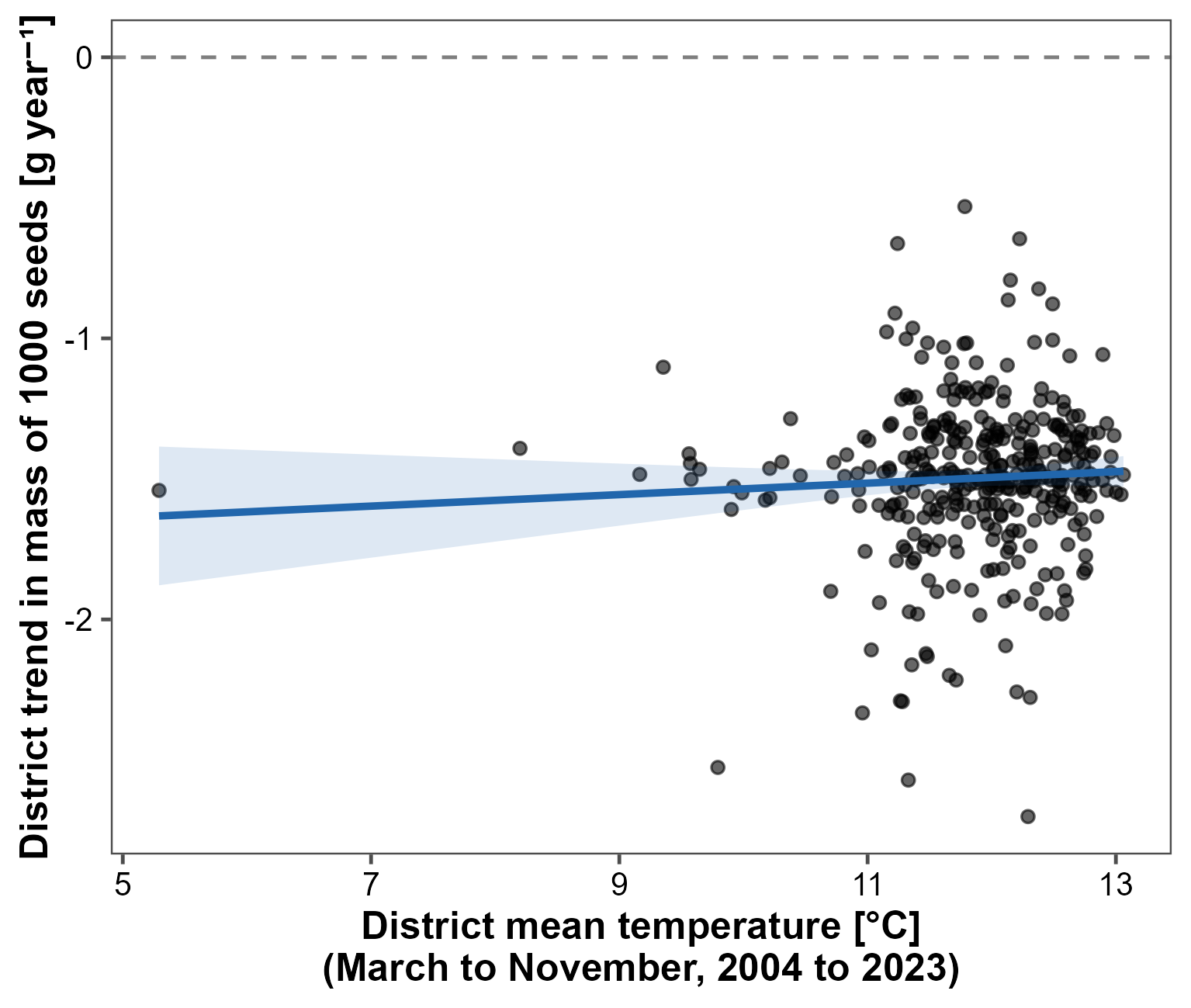

Fig. S3. District-specific temporal trends in the mass of 1000 seeds (conditional slopes from the mixed model with random slopes for year within district) in relation to the mean temperature of the district from March to November 2004 to 2023. The line shows a linear regression with its 95% confidence band; the trends did not depend on district temperature (Table S16). Slopes are shrunk towards the mean and are shown for illustration only. Temperature is given in °C.
